# Chromatin Remodeling by the Histone Methyltransferase SETD2 Drives Lipotoxic Injury in Cardiometabolic Heart Failure with Preserved Ejection Fraction

**DOI:** 10.1101/2024.07.25.605217

**Authors:** Samuele Ambrosini, Sarah Costantino, Shafeeq A. Mohammed, Era Gorica, Melissa Herwig, Loek van Heerebeek, Alessandro Mengozzi, Gergely Karsai, Thorsten Hornemann, Omer Dzemali, Frank Ruschitzka, Nazha Hamdani, Francesco Paneni

## Abstract

**Background:** Cardiometabolic heart failure with preserved ejection fraction (cHFpEF) is highly prevalent and associates with a poor outcome. Pathological gene expression in heart failure is accompanied by changes in active histone marks without major alterations in DNA methylation. Histone 3 trimethylation at lysine 36 (H3k36me3) - a chromatin signature induced by the histone methyltransferase SETD2 - strongly correlates with changes in gene expression in human failing hearts; however, its role is poorly understood. Here we investigate the role of SETD2 in cHFpEF.

**Methods:** Mice with cardiomyocyte-specific deletion of SETD2 (c-SETD2^-/-^) were generated and subjected to high fat diet feeding and L-NAME treatment for 15 weeks to induce cHFpEF. Cardiac function and exercise tolerance were assessed by echocardiography and Treadmill exhaustion test. Chromatin immunoprecipitation assays (ChIP) were performed to investigate SETD2/H3k36me3 enrichment on gene promoters. SETD2 gain- and loss-of-function experiments were performed in cultured cardiomyocytes (CMs) exposed to palmitic acid (PA). SETD2 expression was also investigated in left ventricular (LV) myocardial specimens from patients with cHFpEF and control donors.

**Results:** SETD2 was upregulated in cHFpEF mouse hearts and its chromatin mark H3k36me3 was enriched on the promoter of sterol regulatory element-binding transcription factor 1 (SREBP1) gene. SETD2 activation in cHFpEF led to SREBP1 upregulation, triglyceride accumulation and lipotoxic damage. Of note, cardiomyocyte-specific deletion of SETD2 in mice prevented HFpEF-related hypertrophy, diastolic dysfunction and lung congestion while improving exercise tolerance. SETD2 deletion blunted H3K36me3 enrichment on SREBP1 promoter thus leading to a marked rewiring of the cardiac lipidome and restoration of autophagic flux. SETD2 depletion in PA-treated CMs prevented SREBP1 upregulation, whereas SETD2 overexpression recapitulated lipotoxic damage. Finally, SETD2 was upregulated in LV specimens from cHFpEF patients and its pharmacological inhibition by EZM0414 attenuated CM stiffness.

**Conclusions:** Therapeutic modulation of SETD2/H3k36me3 axis might prevent lipotoxic injury and cardiac dysfunction in cHFpEF.

## Introduction

Heart failure with preserved ejection fraction (HFpEF) is a heterogeneous syndrome presenting with different clinical phenotypes.^1, 2^ Among these, cardiometabolic HFpEF (cHFpEF) is emerging as the most prevalent form of HFpEF.^3, 4^ The pathophysiological mechanisms of cHFpEF remain poorly understood. However, recent advances in the preclinical modeling of the syndrome, coupled with a better definition of its clinical presentations and analysis of human HFpEF myocardial specimens, have unveiled metabolic disturbances and inflammatory burden as key drivers of the cHFpEF pathophysiology.^3^

While genetic mutations account for a small part of the risk of developing cardiovascular disease, environmental cues and lifestyle changes play a central role, especially in a multifactorial syndrome such as cHFpEF.^5^ Epigenetic modifications - defined as plastic changes of chromatin that do not alter DNA sequence - have recently emerged as key players in the pathophysiology of cardiovascular disease.^6, 7^ Post-translational modifications of histones (i.e. methylation, acetylation) are induced by different families of chromatin-modifying enzymes and regulate chromatin accessibility to transcription factors, thus enabling or repressing gene transcription. Recent studies have demonstrated that pathological gene expression in heart failure is mainly accompanied by changes in active histone marks.^8^ Overexpression of Rae28, a protein responsible for H3K27 tri-methylation, led to cardiomyocyte apoptosis, dilated cardiomyopathy, and heart failure in mice.^9^ Similarly, loss of DOT1L, responsible for H3K79me2/3, was associated with downregulation of dystrophin with subsequent development of a cardiomyopathy phenotype.^10^ Although previous work has provided important insights on the role of chromatin modifications in cardiac disease and heart failure, the role of chromatin remodeling in cHFpEF remain elusive. A better understanding of gene-environment interactions and epigenetic changes could help unveiling new mechanisms-based strategies to tackle this multi-systemic syndrome.

Among the constellation of chromatin marks, the histone 3 trimethylation at lysine 36 (H3k36me3) - a chromatin signature specifically induced by the mammalian histone methyltransferase SETD2 – has shown a strong correlation with changes in gene expression in human failing hearts.^11, 12^ H3k36me3 is recognized by several chromatin-modifying enzymes, including histone methyl- and acetyltransferases, DNA methyltransferases, and transcription elongators, making SETD2 a master regulator of chromatin activity and gene transcription in disease states.^13, 14^ To date, only few studies have investigated the role of SETD2 and H3k36me3 in the cardiovascular system. SETD2 was recently found to be involved in hypoxia-induced pulmonary arterial hypertension in mice, and its conditional deletion in vascular smooth muscle cells conferred protection against right ventricular remodeling and dysfunction.^15^ Other studies have shown a role for SETD2 in myogenesis and cardiac development.^16, 17^ The involvement of SETD2/ H3k36me3 axis in the setting of heart failure in general, and HFpEF in particular, is largely unknown.

Here we show that SETD2 is upregulated in the cHFpEF myocardium (specifically in cardiomyocytes) and governs the transcription of lipotoxic genes (i.e. SREBP1) fostering lipogenic transcriptional programs with subsequent accumulation of triglycerides and ceramides, impaired autophagic flux and cardiac dysfunction. Notably, cardiomyocyte-specific deletion of SETD2 in mice blunted SREBP1 signaling thus rewiring the myocardial lipidome and rescuing cHFpEF-related cardiac remodeling and dysfunction. Of clinical relevance, we show that selective pharmacological inhibition of SETD2 in skinned CMs obtained from LV specimens of cHFpEF patients was able to attenuate CM passive stiffness, a major pathological feature of HFpEF.

## Material and Methods

A detailed description of the experiments in cultured cardiomyocytes, cardiomyocyte-specific SETD2 knockout mice (*α-MHC–SETD2^-/-^*) and LV human specimens is reported in detail in the Supplemental Material. All the primers used for real time PCR are reported in **Table S1.**

### Cardiac-specific SETD2 knockout mice (α-MHC–SETD2^-/-^)

SETD2^fl/+^ (C57BL/6N-Setd2^<tm1c(NCOM)Mfgc>^Tcp) mice were obtained from the Canadian Mouse Mutant Repository, Hospital for Sick Children, Canada and cross bred to generate SETD2^fl/fl^. SETD2^fl/fl^ were bred with α-MHC-Cre mice to generate cardiomyocyte-specific (c-SETD2^-/-^) knockout animals.

### Mouse model of cardiometabolic HFpEF

To develop cHFpEF, both c-SETD2^-/-^ and SETD2^fl/fl^ underwent 15 weeks of HFD (60 kcal% fat, D12492) plus L-NAME (0.5 g/liter in the drinking water) or normal diet (10 kcal% fat, D12450J) in the control group, as previously reported.^18^ Animal experiments were approved by the Kommission für Tierversuche des Kantons Zürich, Switzerland.

### SETD2 inhibition in skinned cardiomyocytes from patients with cHFpEF

Force measurements were performed on single de-membranated cardiomyocytes isolated from left ventricular specimens obtained from patients with cHFpEF and control donors for heart transplant (2–3 cardiomyocytes per biopsy) collected at the at the Department of Cardiology, OLVG, Amsterdam, the Netherlands, as described previously.^19^ Skinned CMs were incubated with the selective SETD2 inhibitor EZM0414 (10μM) or vehicle for 60 min followed by assessment of passive myofilament stiffness. All procedures were approved by the local ethics committee and performed in accordance with the Declaration of Helsinki. A detailed description of the study population is reported in Supplemental Material.

### Statistical analysis

The normality of continuous variables was assessed by the Kolmogorov–Smirnov test. All normally distributed variables are expressed as mean ±standard deviation, unless otherwise stated. Comparisons of continuous variables were performed using unpaired two-sample t-test and Mann–Whitney U test, as appropriate. Categorical variables were compared by using the χ2 test. Multiple comparisons between normally distributed variables were performed by one-way analysis of variance (ANOVA), followed by Bonferroni correction or Benjamini and Hochberg Discovery Rate (FDR). The latter was used when the n number was equal to or lower than the number of experimental groups. Variable correlations were assessed by Spearman’s test.

Probability values <0.05 were considered statistically significant. All analyses were performed with GraphPad Prism Software (version 7.03).

## Results

### Upregulation of SETD2/H3K36me3 associates with lipotoxic damage in cardiometabolic HFpEF

ChIP-Seq datasets (GEO: GSE107785) were interrogated to appraise the role of the active mark H3k36me3 in regulating gene expression. We found that H3k36me3 was involved in the transcriptional regulation of several signaling pathways, including cellular metabolism, protein transport, and degradation. Among top-ranking pathways shown by gene ontology (GO) analysis, the Sterol Regulatory Element-Binding Protein (SREBP) signaling was found to be of particular interest in the context of cardiometabolic alterations, given its central role in the modulation of lipid biosynthesis and uptake (**Fig. 1A, Table S2**). We also found that H3k36me3 was enriched on SREBP1 gene promoter and clustered with active regulatory marks such H3k4me3 and H3k27ac (GEO: GSE75270, **Fig 1B**). To investigate the role of SETD2/H3k36me3 signaling in cHFpEF, we employed a validated two-hit mouse model of HFpEF, which combines metabolic and hemodynamic stress and has been shown to faithfully resemble features of human cHFpEF.^18, 20^ We observed that SETD2 protein expression was significantly increased in cHFpEF mouse hearts as compared to controls and was associated with increased H3k36me3 levels (**Fig. 1C**). To detect cell-specific changes of SETD2 expression within the cHFpEF myocardium, we next investigated SETD2 expression pattern in different cardiac cells, namely cardiomyocytes, endothelial cells, fibroblasts and immune cells. We found that SETD2 was preferentially upregulated in CMs from cHFpEF mice (**Fig. S1**). ChIP assays performed in mouse cardiomyocytes from cHFpEF and control mice confirmed the enrichment of H3k36me3 on *Srebp1* gene promoter and enhanced *Srebp1* transcription, as shown by real-time PCR (**Fig 1D**). Protein-protein interaction (PPI) analysis showed that SREBP1 interacts with molecular partners involved in *de novo* lipid synthesis and generation of mono-unsaturated fatty acids, such as fatty acid synthase (FASN), stearoyl-CoA desaturase-1 (SCD1) and acetyl-CoA carboxylase 1 (ACC1) (**Fig. 1E**). We found that *Fasn*, *Scd1* and *Acc1* were upregulated in cHFpEF hearts as compared to controls (**Fig. 1F**). In line with these results, SREBP1 upregulation in the HFpEF myocardium was associated with accumulation of triglycerides (TG) in the heart (**Fig. 1G**). Excess fat accumulation and lipotoxic stress have recently been shown to promote cardiomyocyte damage by suppressing autophagy.^21–24^ In cHFpEF hearts, we found that *mTOR* was upregulated and was associated with increased activation of the autophagy inhibitor mTORC1, as assessed by phosphorylation of ribosomal protein S6 kinase beta (S6K) (**Fig. 1H**).^25^ Consistently, autophagy-related genes *Atg7* and *Atg13* and autophagic flux were suppressed in the HFpEF myocardium (**Fig. 1I-J**) while myocardial apoptosis was enhanced (**Fig. 1K**).

**Figure 1.**
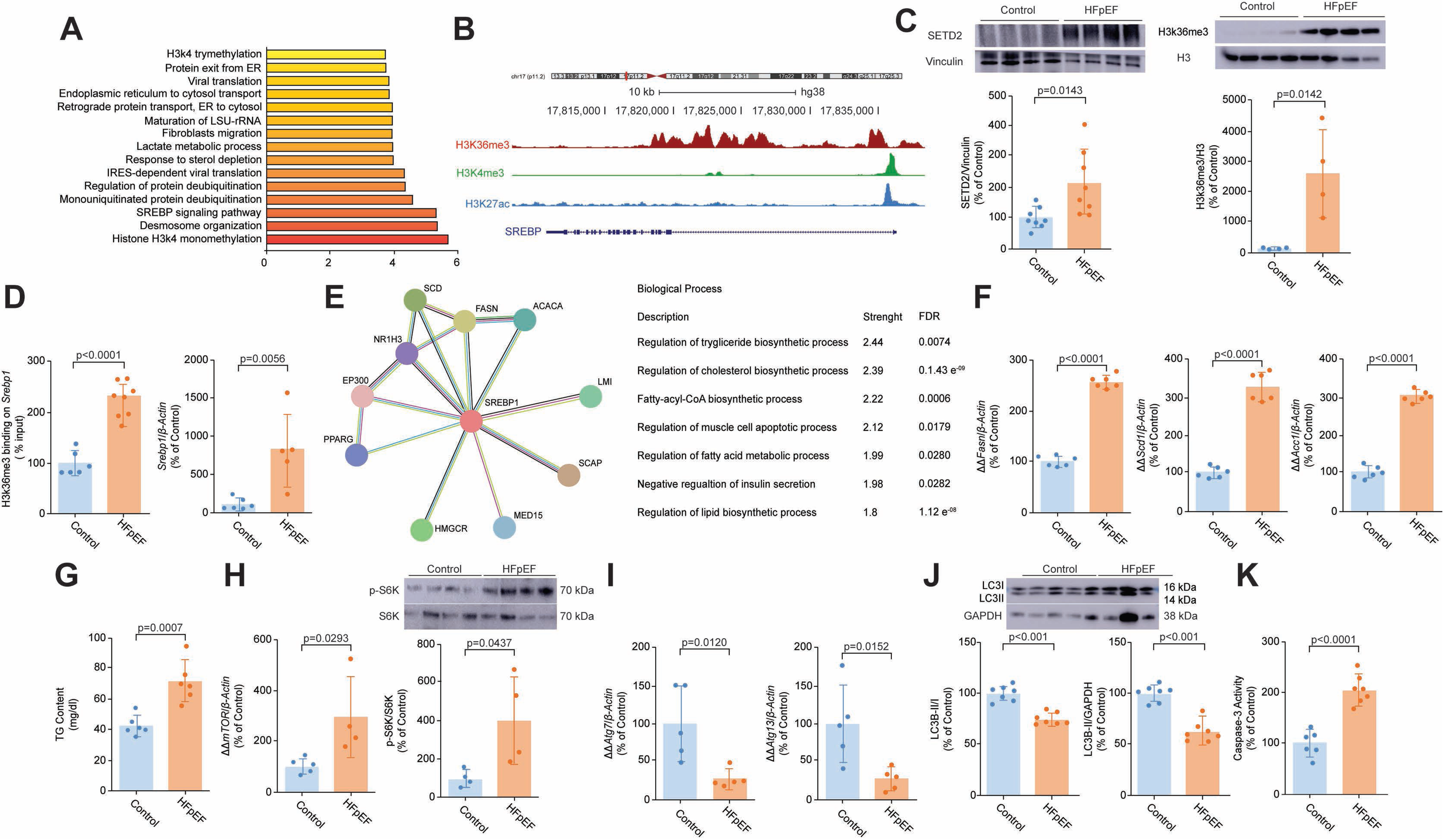
SETD2/H3K36me3 activation and lipotoxic damage in cardiometabolic HFpEF. **A**) ChIP-seq data showing pathway enrichment analysis for the H3k36me3 chromatin mark (GEO: GSE107785); **B**) Bioinformatic analysis of ChIP-seq data (GEO: GSE75270) showing enrichment of H3k36me3 and other chromatin activating marks (H3k27ac, H3k4me3) on the *Srebp1* promoter; **C**) Western blot and relative quantification showing protein expression of SETD2 and H3k36me3 in LV specimens from HFpEF and control mice; **D**) ChIP assay showing H3k36me3 enrichment on *Srebp1* promoter and *Srebp1* gene expression in HFpEF and control mouse hearts. Chromatin was immunoprecipitated with an antibody against H3k36me3 and precipitated genomic DNA was analyzed by real-time PCR using primers for *Srebp1* promoter; **E**) Protein-protein interaction (PPI) analysis showing SREBP1-related targets and top-ranked biological processes; **F**) Real time PCR showing expression of SREBP1-dependent genes *Fasn*, *Scd1* and *Acc1;* **G**) Triglyceride content in HFpEF and control hearts; **H**) Real time PCR showing gene expression of mTOR. Western blot and relative quantification show phosphorylated S6K, a marker of mTORC1 activity; **I**) Gene expression of the autophagosome markers *Atg7* and *Atg13* in HFpEF and control hearts; **J**) Western blot and relative quantification showing LC3II/I ratio and LC3B-II protein expression in HFpEF and control hearts; **K**) Apoptosis assessed by Caspase-3 Assay in LV specimens from controls and HFpEF mice. Results are presented as mean±SD. Group comparisons were performed by Student’s t-test. A p-value <0.05 versus control was considered significant. HFpEF: Heart failure with preserved ejection fraction, Srebp1: Sterol regulating element-binding protein 1, TG: Triglycerides, mTOR: Mechanistic Target of Rapamycin, S6K: Ribosomal protein S6 kinase beta, Atg: Autophagy related gene, Fasn: Fatty Acid Synthase, Scd1: Stearoyl-CoA desaturase-1, Acc1: Acetyl-CoA carboxylase 1.

### Cardiomyocyte-specific deletion of SETD2 prevents cHFpEF development in mice

To appraise the *in vivo* role of SETD2 in cHFpEF, we generated cardiomyocyte (CM)-specific SETD2 knockout mice (c-SETD2^-/-^) and used SETD2^fl/fl^ littermates as a control group. Mice lacking SETD2 in CMs had significantly lower levels of H3k36me3 in the heart but not in other tissues (**Fig. S2**). At the age of 8 weeks, c-SETD2^-/-^ and SETD2^fl/fl^ mice were fed HFD and chronically treated with L-NAME for 15 weeks to induce cHFpEF. Control groups were fed a normal chow diet. Compared to SETD2^fl/fl^ littermates, c-SETD2^-/-^ mice did not display alterations in body weight in either control or cHFpEF groups (**Fig. 2A**). SETD2^fl/fl^ cHFpEF mice displayed cardiac hypertrophy and lung congestion, key features of HFpEF, whereas these changes were not observed in c-SETD2^-/-^ cHFpEF animals (**Fig 2B-C**). Systolic function - assessed by left ventricular ejection fraction - was preserved and comparable among the different experimental groups (**Fig. 2D**). Echocardiography showed that SETD2^fl/fl^ cHFpEF mice had increased left ventricular mass (LVM) and diastolic dysfunction - assessed by E/A ratio, E/e’ ratio and isovolumic relaxation time (IVRT) - whereas LV mass and diastolic function were preserved in cHFpEF mice lacking SETD2 (**Fig 2E-G, Fig. S3-5**). Consistently, myocardial performance index (MPI) – a well-established index of myocardial function and a predictor of heart failure in humans^26^ - was impaired in SETD2^fl/fl^ cHFpEF mice but preserved in c-SETD2^-/-^ cHFpEF animals (**Fig 2H, Fig. S5**). Moreover, exercise tolerance assessed by Treadmill exhaustion test was impaired in SETD2^fl/fl^ HFpEF mice, while an improvement was observed in mice with CM-specific deletion of SETD2 (**Fig 2I**).

**Figure 2.**
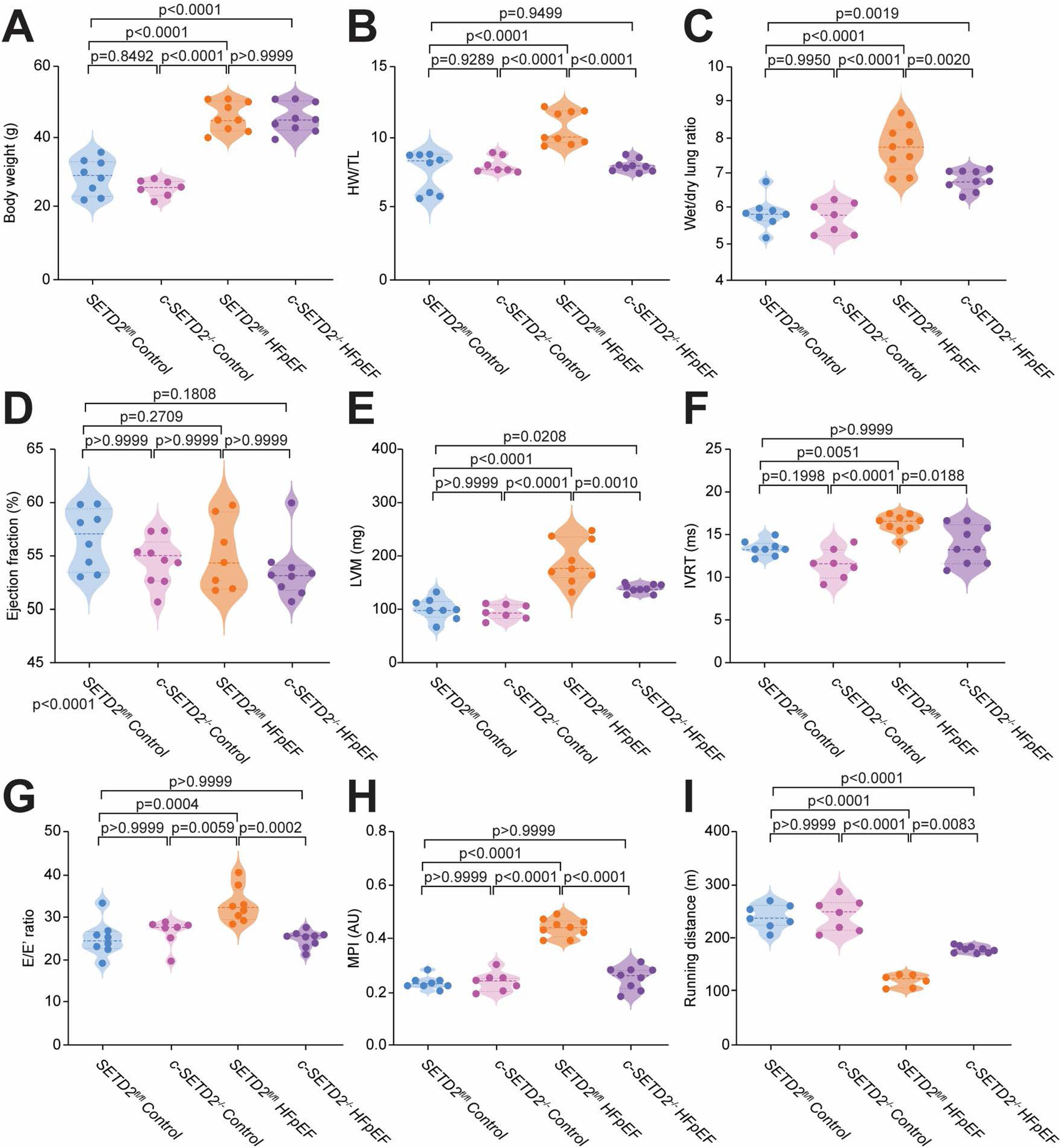
Cardiomyocyte-specific deletion of SETD2 protects against cardiometabolic HFpEF. **A**) Body weight in HFpEF and control mice, with and without CM-specific deletion of SETD2; **B-C**) Heart weight/tibia length and lung wet/dry ratio across the 4 experimental groups; **D**) Systolic function assessed by left ventricular ejection fraction (EF); **E**) Left ventricular mass assessed by echocardiography in the four experimental groups; **F-G**) Diastolic function as assessed by IVRT and E/A ratio; **H**) Myocardial performance index; **I**) Exercise tolerance in the 4 experimental groups, assessed as running distance by Treadmill exhaustion test. Results are presented as means ± SD and compared by one-way analysis of variance (ANOVA) followed by post-hoc analysis with Bonferroni correction. Adjusted p values (alpha/g) are shown. A p-value <0.05 was considered significant. HW/TL: Heart weight/tibia length ratio, LVM: Left ventricular mass, IVRT: Isovolumic relaxation time; MPI: Myocardial performance index.

### SETD2 drives HFpEF-related lipotoxic damage and defective autophagy in vivo

Given the specificity of SETD2 for H3k36me3, we next checked the expression of this chromatin mark in mice lacking SETD2. H3k36me3 levels were increased in cHFpEF SETD2^fl/fl^ hearts as compared to controls but significantly reduced in c-SETD2^-/-^ groups (**Fig. 3A**). ChIP assays revealed the enrichment of H3k36me3 on the promoter of *srebp1* in SETD2^fl/fl^ cHFpEF mice, while no enrichment was found in mice lacking SETD2 (**Fig. 3B**). Furthermore, *Srebp1* was upregulated in SETD2^fl/fl^ cHFpEF animals compared to controls, and its expression was reduced in c-SETD2^-/-^ mice (**Fig. 3C**), suggesting that SETD2 is required for *Srebp1* transcription. Along the same line, *Fasn*, *Scd1* and *Acc1* - SREBP1-dependent genes involved in lipid metabolism - were upregulated in the SETD2^fl/fl^ cHFpEF group but not in mice lacking SETD2 (**Fig. 3D**). Given the involvement of SETD2 in the transcriptional control of lipid genes, we next investigated intra-myocardial TG content in the different experimental groups.

**Figure 3.**
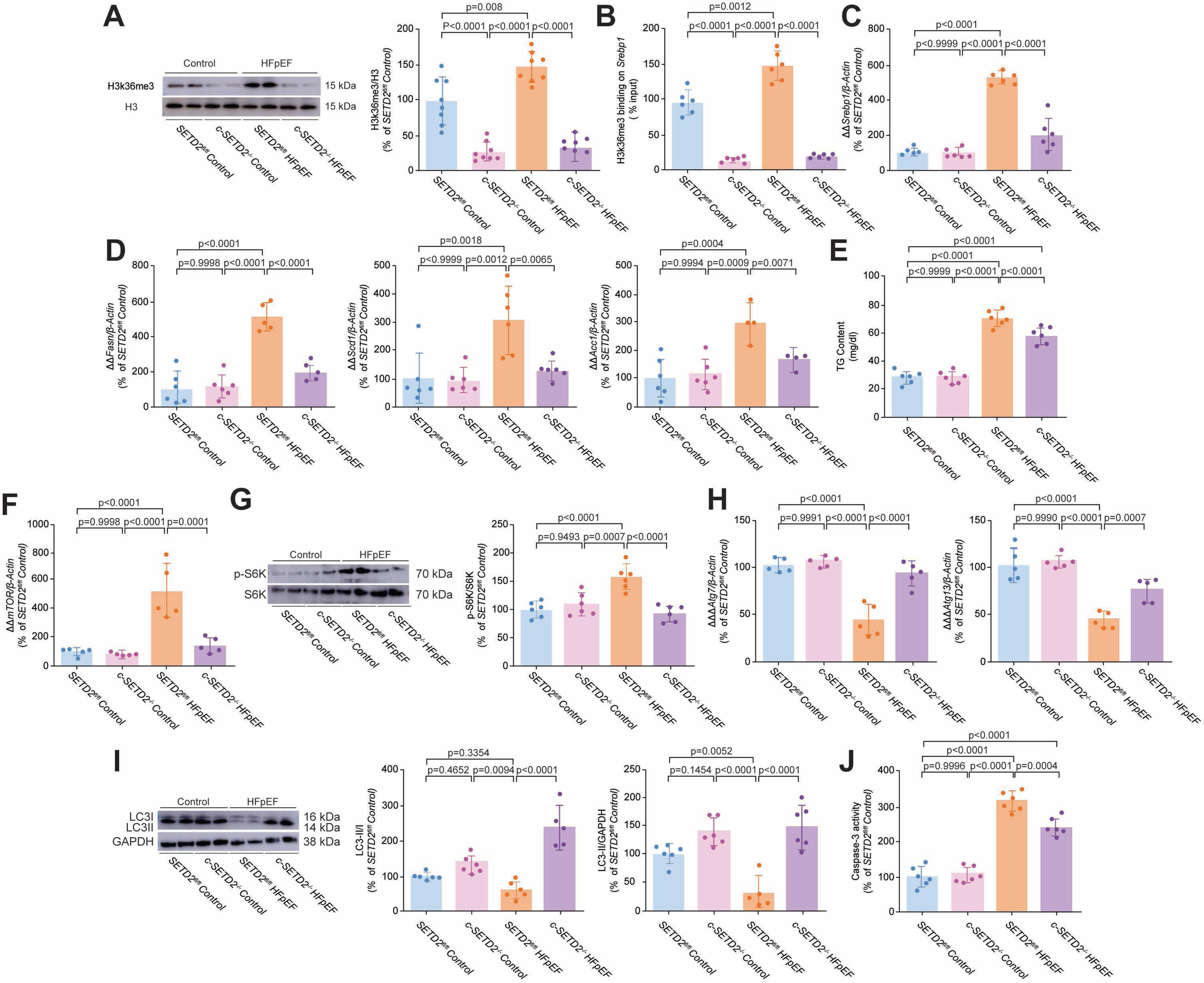
Lipotoxic injury and defective autophagy are prevented by SETD2 deletion in HFpEF mice. **A**) Western blot (representative images and relative quantification) showing expression of H3k36me3 in HFpEF and control mouse hearts, in the presence or in the absence of cardiomyocyte-specific SETD2 deletion; **B**) ChIP assay showing H3k36me3 enrichment on the SREBP1 promoter in LV specimens from the 4 experimental groups. Chromatin was immunoprecipitated with an antibody against H3k36me3 and precipitated genomic DNA was analyzed by real-time PCR using primers for *Srebp1* promoter; **C-D**) Real-time PCR showing gene expression of *Srebp1* and its target genes *Fasn*, *Scd1* and *Acc1*. **E**) Intra-myocardial triglyceride content across the 4 experimental groups; **F**) Real time PCR showing gene expression of mTOR; **G**) Western blot and relative quantification show phosphorylated S6K, a marker of mTORC1 activity; **H**) Gene expression of autophagy-related genes *Atg7* and *Atg13*; **I**) Autophagic flux assessed by Western blot of LC3II/I and relative quantification; **J**) Myocardial apoptosis assessed by Caspase-3 activity assay. Results are presented as mean±SD and compared by one-way analysis of variance (ANOVA) followed post-hoc analysis with Bonferroni correction. Adjusted p values (alpha/g) are shown. A p-value <0.05 was considered significant. HFpEF: Heart failure with preserved ejection fraction, Srebp1: Sterol regulating element-binding protein 1, TG: Triglycerides, mTOR: Mechanistic Target of Rapamycin, S6K: Ribosomal protein S6 kinase beta, Atg: Autophagy related gene. Fasn: Fatty Acid Synthase, Scd1: Stearoyl-CoA desaturase-1, Acc1: Acetyl-CoA carboxylase 1.

Consistent with our observations, total TG levels were increased in LV specimens from SETD2^fl/fl^ cHFpEF as compared with control mice (**Fig. 3E**). In contrast, intramyocardial TG levels were reduced in mice with conditional deletion of SETD2 as compared to SETD2^fl/fl^ cHFpEF littermates (**Fig. 3E**). Lipid overload in SETD2^fl/fl^ cHFpEF mice was associated with impaired autophagy, as shown by enhanced *mTOR* expression, mTORC1 activation and increased apoptosis, as assessed by Caspase-3 activity (**Fig 3G-J**). By contrast, mTOR signaling, autophagic flux and apoptosis were preserved in mice lacking SETD2 (**Fig. 3G-J**).

### SETD2 deletion rewires the cardiac lipidome in cHFpEF mice

To investigate the full spectrum of lipid species in our setting, we performed MS-based lipidomic analysis in LV samples from control and cHFpEF mice, with and without SETD2 deletion. In cHFpEF mice, we observed profound changes of several lipid species (including several subclasses of TGs) as compared to control animals (all the raw data and statistical comparisons for cardiac lipidomics are reported in the **Appendix**). Of interest, SETD2 deletion in CMs was associated with an extensive lipidome rewiring, with a near normalization of almost all TG subclasses, namely medium-long chain TGs, long chain TGs and very long chain TGs (**Fig. 4A-B**). A predominant effect on the first two subclasses was observed, suggesting a role for SETD2 in modulating *de novo* synthesis of fatty acids (**Fig. 4A-B**). Besides alterations in TGs levels, we observed a remarkable increase in ceramides levels, namely Cer(d18:1/18:0) and Cer(d18:1/24:0) in the HFpEF myocardium as compared to control mice (**Fig. 4C-D**). Of interest, these changes were not observed in mice lacking SETD2. By contrast, lipidomic analysis performed in mouse plasma did not show relevant changes between cHFpEF mice with and without SETD2 deletion, suggesting that cardiac-specific lipid signals drive lipotoxic injury in cHFpEF (**Fig. S6**).

**Figure 4.**
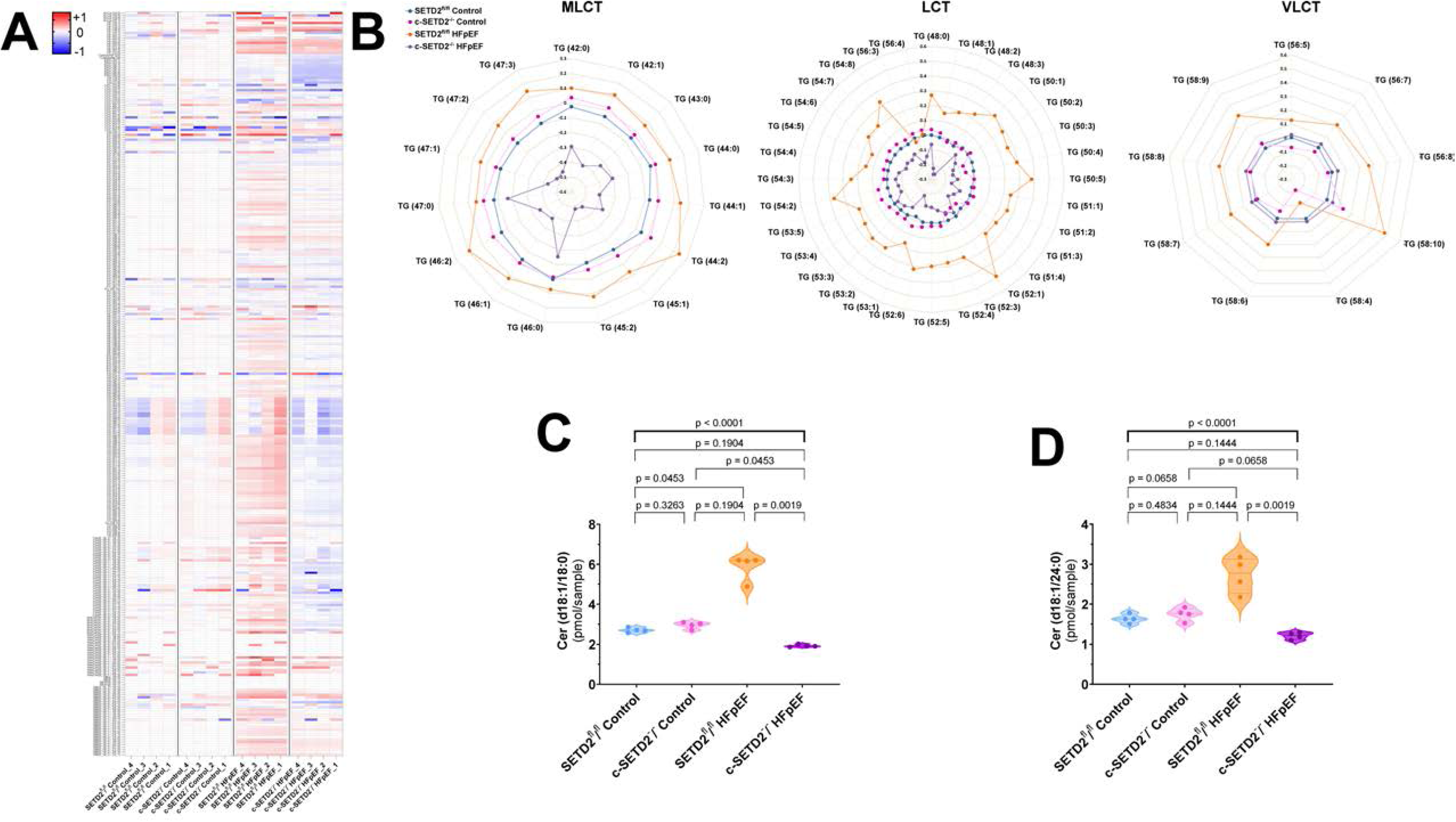
SETD2 deletion rewires the myocardial lipidome in HFpEF mice. **A**) Heat map showing changes of the myocardial lipidome across the 4 experimental groups (n=4/group); **B**) Radar plots showing cardiac levels of medium, long-chain and very long-chain TGs in the 4 experimental groups (n=4/group). Data are presented as median value and expressed in log10 changes from controls; each grey line represents a 0.1-fold change; **C-D**) Cardiac levels of Cer(d18:1/18:0)] and [Cer(d18:1/24:0)] across the 4 experimental groups. LCT: long-chain triacylglycerols; MLCT: medium and long-chain triacylglycerols; ND: normal diet; TGs: triglycerides; VLCT: very long-chain triacylglycerols.

### Metabolic stress induces SETD2/H3k36me3 dysregulation and lipotoxic injury in cardiomyocytes

To further investigate SETD2 signaling in cardiomyocytes, we performed *in vitro* experiments in cultured CMs exposed to palmitic acid (PA), a well-established model mimicking metabolic stress and lipotoxic damage.^27^ Consistent with our in vivo observations, PA-treated CMs displayed increased H3k36me3 levels, enhanced H3k36me3 enrichment on *Srebp1* promoter, and subsequent increase of *Srebp1* transcription (**Fig. 5A-B**). Upregulation of SREBP1 was associated with TG accumulation and suppression of autophagy (**Fig. 5C-F**). Lipotoxic injury in our experimental setting was also outlined by increased Caspase-3 activation and decreased cell survival (**Fig. 5G**).

**Figure 5.**
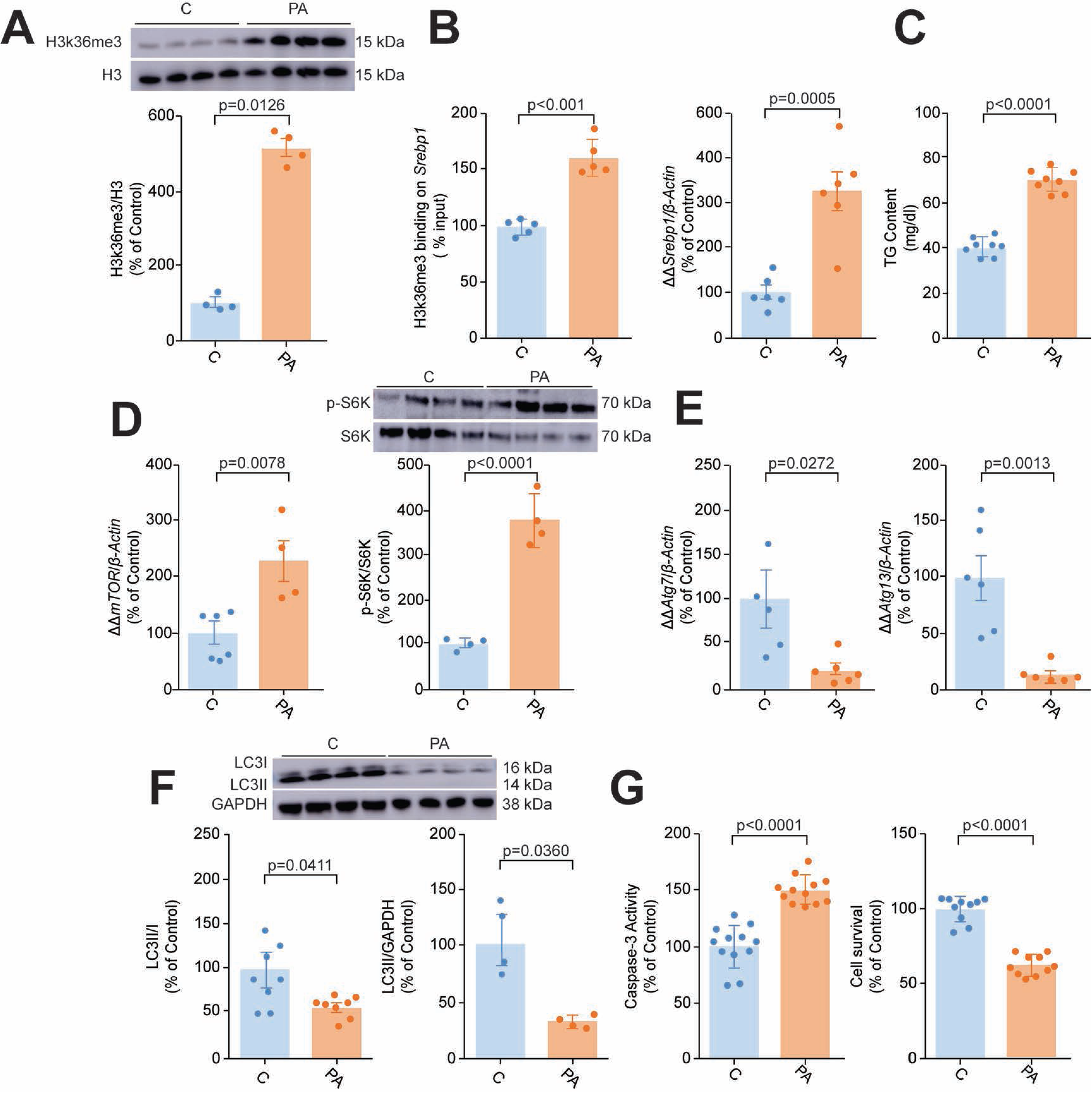
Metabolic stress induces SETD2/H3k36me3 deregulation and lipotoxic injury in cardiomyocytes. **A**) Western blot (representative images and relative quantification) showing expression H3k36me3 in H9c2 cells treated with palmitic acid (PA) or control for 48h; **B**) ChIP assay showing the enrichment of H3k36me3 on the *srebp1* promoter in PA-treated cells and control; **C**) Triglyceride content in H9c2 cells exposed to PA or control; **D**) Real time PCR showing gene expression of mTOR. Western blot and relative quantification show phosphorylated S6K, a marker of mTORC1 activity; **E**) Gene expression of *Atg7* and *Atg13*; **F**) Autophagic flux assessed by Western blot of LC3II/I and relative quantification in PA-treated cells and control; **G**) Cardiomyocyte apoptosis and cell survival, assessed by Caspase-3 activity and CellTiter Blue assay, respectively. Results are presented as means ± SD. A p-value <0.05 versus C by Student’s t-test was considered significant. C: control, PA: Palmitic acid, Srebp1: Sterol regulating element-binding protein 1, TG: Triglycerides, mTOR: Mechanistic Target of Rapamycin, S6K: Ribosomal protein S6 kinase beta, Atg: Autophagy related gene.

### SETD2 depletion prevents lipotoxic damage while preserving autophagy

We next asked whether SETD2 inhibition could prevent transcriptional programs involved in lipotoxic injury. SiRNA-mediated knockdown of SETD2 led to a significant reduction of *Setd2* expression and blunted H3k36me3 levels, thus confirming SETD2 specificity for this chromatin signature (**Fig. 6A, Fig. S7**). SETD2 depletion in CMs led to reduced H3k36me3 enrichment on *Srebp1* promoter, thus preventing palmitic acid-induced SREBP1 upregulation and TG accumulation (**Fig. 6B-C**). Reduced TG uptake following SETD2 knockdown was associated with suppression of *mTOR* expression and activity, upregulation of autophagy-related genes, and restoration of the autophagic flux (**Fig. 6D-F**). Moreover, the prevention of lipotoxic damage upon SETD2 inhibition was also demonstrated by reduced apoptosis and improved cell survival (**Fig. 6G**).

**Figure 6.**
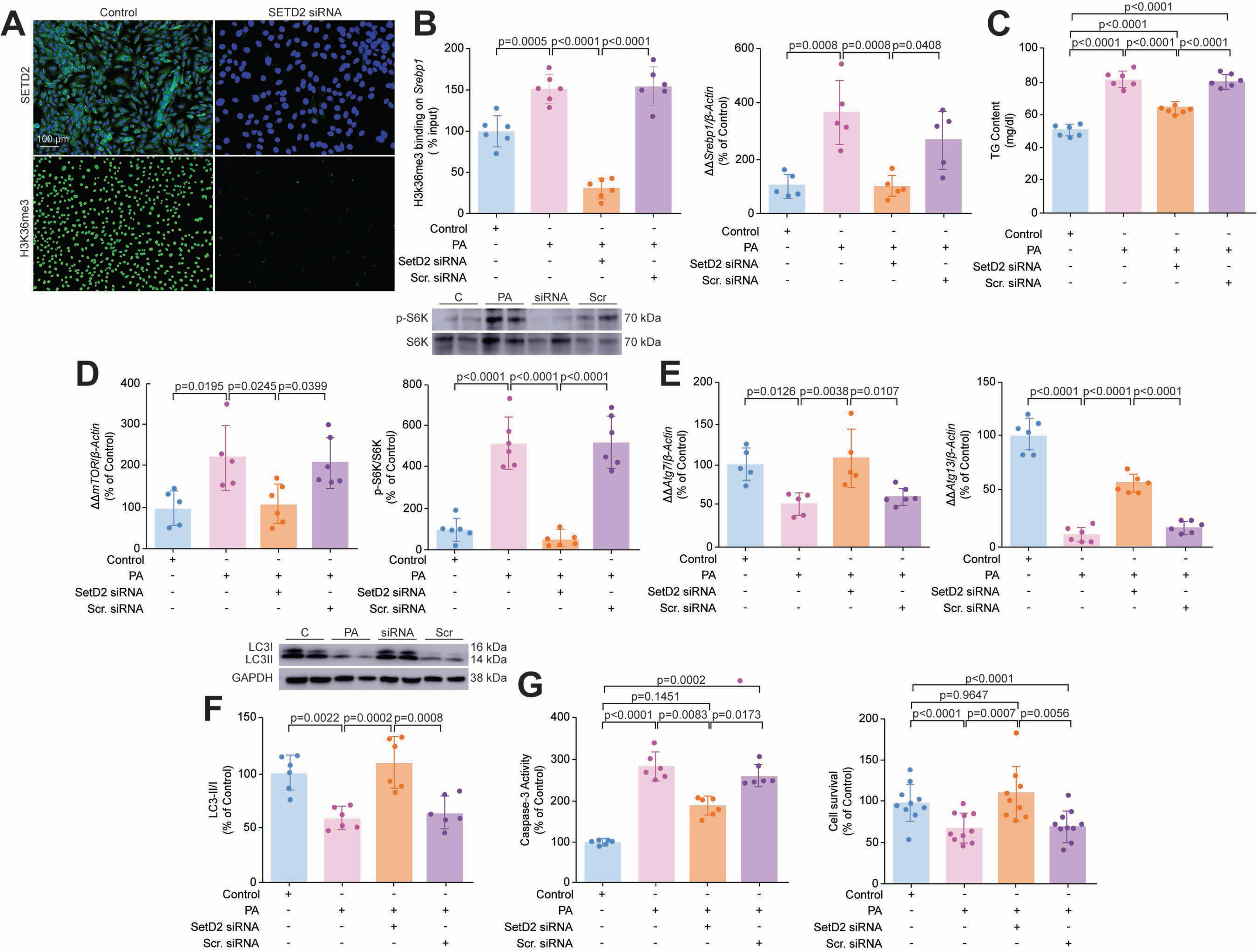
SETD2 depletion blunts H3K36m3 levels, SREBP1 expression, and lipotoxic damage in cultured cardiomyocytes. **A**) Immunofluorescence showing siRNA-mediated SETD2 knockdown and its effect on H3k36me3 levels (green). Nuclei are stained with DAPI (blue). Images are representative of 4 independent experiments; **B**) ChIP assay showing H3k36me3 enrichment on the *srebp1* promoter in control and PA-treated CMs, in the presence of SETD2 siRNA or Scr.siRNA; **C**) Triglyceride content in control and PA-treated CMs, in the presence of SETD2 siRNA or Scr.siRNA; **D**) Real time PCR showing gene expression of mTOR. Western blot and relative quantification show phosphorylated S6K, a marker of mTORC1 activity; **E**) Gene expression of autophagy-related genes *Atg7* and *Atg13*; **F**) Autophagic flux assessed by Western blot and relative quantification of LC3II/I in control and PA-treated cells, in the presence of SETD2 siRNA or Scr.siRNA; **G**) Cardiomyocyte apoptosis and cell survival, assessed by Caspase-3 activity and CellTiter Blue assay, respectively. Results are presented as mean ± SD and compared by one-way analysis of variance (ANOVA) followed by post-hoc analysis with Bonferroni correction. Adjusted p values (alpha/g) are shown. A p-value <0.05 was considered significant. C: control, PA: Palmitic acid, Srebp1: Sterol regulating element-binding protein 1, TG: Triglycerides, mTOR: Mechanistic Target of Rapamycin, S6K: Ribosomal protein S6 kinase beta, Atg: Autophagy related gene.

### SETD2 overexpression recapitulates lipotoxic injury in cardiomyocytes

To investigate whether SETD2 activation in CMs mimics lipotoxic damage, we performed gain-of-function experiments by overexpressing SETD2. SETD2 overexpression led to a significant increase in H3k36me3 (**Fig. 7A-B**) with subsequent SREBP1 upregulation, TG accumulation and increased apoptosis (**Fig. 7C-F**). To investigate whether SREBP1 is required for the detrimental effects of SETD2 in CMs, we overexpressed SETD2 in the presence of concomitant SREBP1 knockdown. Interestingly, SETD2 overexpression recapitulated lipotoxic injury while concomitant inhibition of SREBP1 prevented lipid accumulation, lipotoxic damage and autophagy impairment (**Fig. 7G-I**).

**Figure 7.**
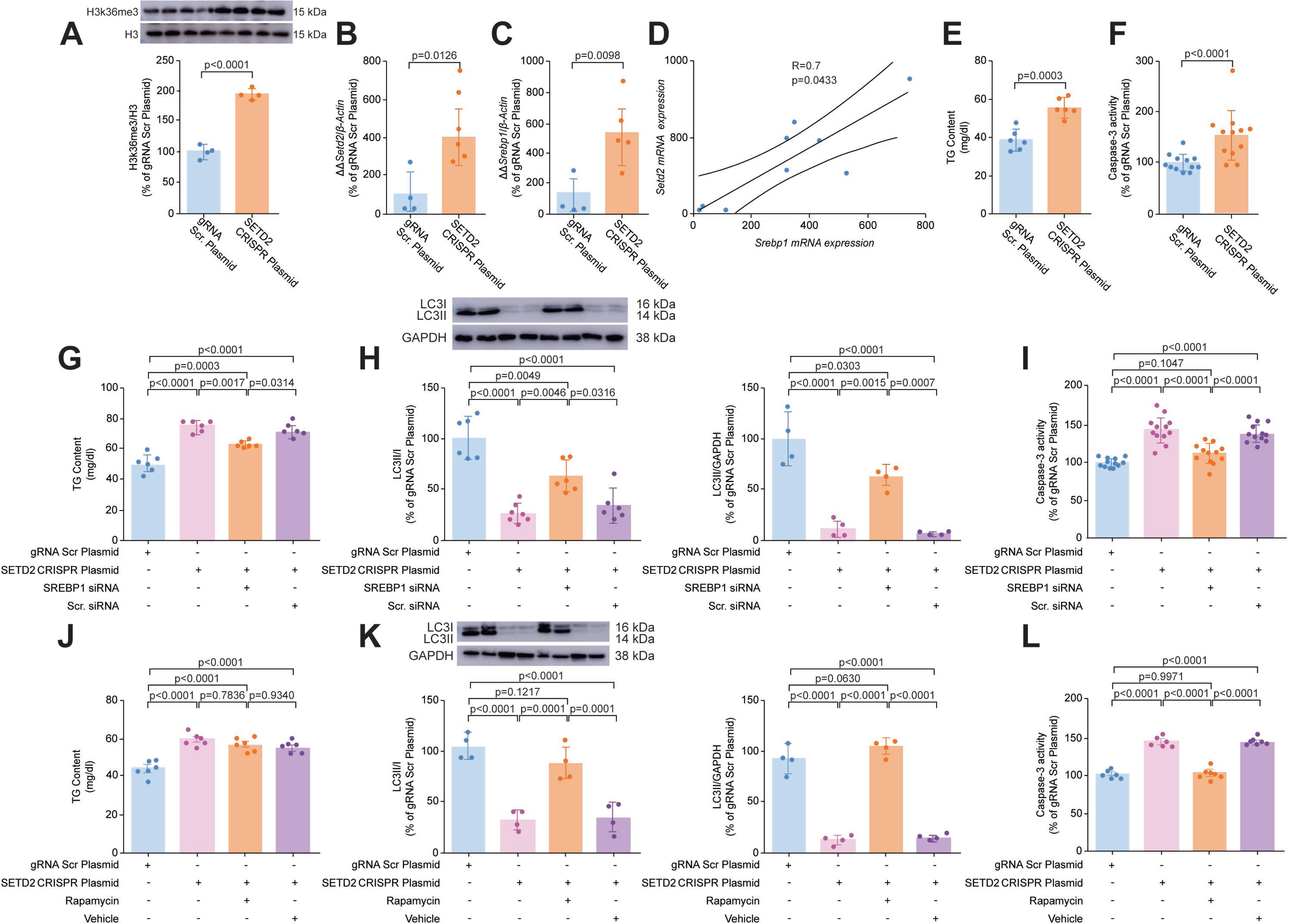
SREBP1 is required for SETD2-dependent lipotoxic damage and impairment of autophagy. **A**) Real time PCR showing SETD2 gene expression in H9c2 cells treated with SETD2 CRISPR Plasmid or Scr. Plasmid; **B)** Western blot (representative images and relative quantification) of H3k36me3 following SETD2 overexpression in H9c2 cells; **C**) real time PCR showing *srebp1* gene upregulation in SETD2-overexpressing cells; **D**) Spearman analysis showing a positive correlation between *Setd2* and *Srebp1* gene expression in H9c2 cells; **E**) Triglyceride content in H9c2 cells with SETD2 overexpression or Scr. Plasmid; **F-G**) Caspase-3 activity in the 2 experimental groups; **G-I**) Effect of SREBP1 depletion in SETD2 overexpressing cells on triglyceride content, autophagic flux (assessed by Western blot of LC3II/I and relative quantification) and apoptosis (assessed by Caspase-3 activity); **J**) Triglyceride content in SETD2 overexpressing CMs, in the presence or in the absence of the autophagy activator Rapamycin; **K-L**) Treatment with Rapamycin rescues autophagic flux (assessed by Western blot of LC3II/I and relative quantification) and apoptosis (assessed by Caspase-3 activity) in SETD2 overexpressing CMs. Results are presented as means ± SD. A p-value <0.05 by one-way analysis of variance (ANOVA) followed by post-hoc analysis with Bonferroni correction was considered significant. Adjusted p values (alpha/g) are shown. C: control, Srebp1: Sterol regulating element-binding protein 1, TGs: Triglycerides.

### Pharmacological reactivation of autophagy rescues lipotoxic injury in CMs

Our in vivo and in vitro data showed that SETD2 activation suppresses autophagy by fostering lipid overload in CMs. Whether defective autophagy is causally implicated in SETD2-mediated lipotoxic injury remains elusive. To appraise the contribution of autophagy in our setting, we treated SETD2-overexpressing CMs with Rapamycin, a well-established pharmacological activator of autophagy (**Fig. S8**). SETD2 overexpression led to TG accumulation, a process that was not reverted by Rapamycin (**Fig. 7J**). However, restoration of autophagy prevented apoptosis in SETD2 overexpressing CMs (**Fig 7K-L**).

### SETD2 upregulation in LV myocardial specimens from cHFpEF patients

To investigate whether our findings hold true in the human HFpEF myocardium, we analyzed left ventricular (LV) samples from patients with cHFpEF and age-matched control donors. We found that SETD2 was upregulated in LV specimens from cHFpEF patients as compared to control donors (**Fig 8A**). Furthermore, we observed that the passive sarcomere length-tension relationship, a marker of cardiomyocyte stiffness^28^, was steeper in cHFpEF patients than controls, and showed a positive correlation with *Setd2* gene expression (**Fig. 8B**).

**Figure 8:**
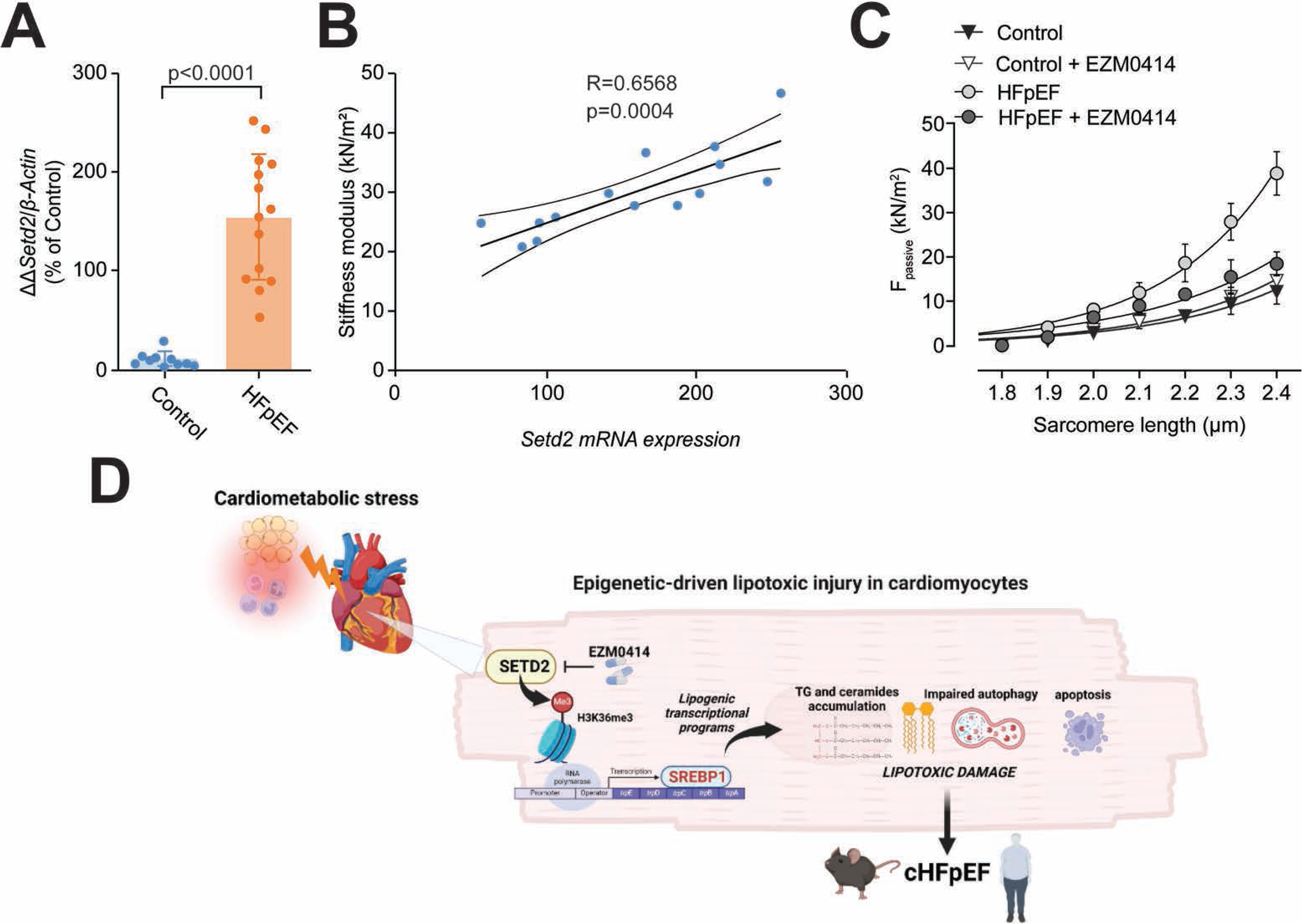
Pharmacological inhibition of SETD2 attenuates passive stiffness in cardiomyocytes from HFpEF patients. **A**) Gene expression of *Setd2* in LV specimens collected from HFpEF patients and age-matched control donors; **B**) Spearman correlation of SETD2 expression with passive stiffness of skinned cardiomyocytes isolated from patients with HFpEF and control donors; **C**) Effects of the selective SETD2 pharmacological inhibitor EZM0414 on passive stiffness in skinned cardiomyocytes isolated from LV specimens of HFpEF patients and age-matched control donors. Results are presented as mean±SD. A p-value <0.05 versus control group by Student’s t-test was considered significant. **D**) Schematic summarizing the role of SETD2 in cardiometabolic HFpEF. In conditions of metabolic stress, activation of SETD2 leads to H3K36me3 on *srebp1* promoter thus promoting gene transcription and protein upregulation. Activation of SREBP1 signaling promotes lipid-related transcriptional programs eventually leading to accumulation of TGs, impairment of autophagic flux and cardiac dysfunction in vivo. Cardiomyocyte-specific deletion of SETD2 was able to blunt H3K36me3 enrichment on *srebp1* promoter thus preventing lipotoxic injury and LV dysfunction in HFpEF mice. HFpEF: Heart failure with preserved ejection fraction.

### Pharmacological inhibition of SETD2 by EZM0414 rescues CM stiffness in patients with cHFpEF

To investigate the potential clinical relevance of SETD2-targeting approaches in human HFpEF, we tested the effects of the selective SETD2 inhibitor EZM0414 in skinned CMs from patients with cHFpEF and control donors. The cardiomyocyte F_passive_ was significantly higher in cHFpEF patients as compared to controls. Of interest, treatment with EZM0414 significantly attenuated cardiomyocyte F_passive_ (**Fig. 8C**).

## Discussion

Our study demonstrates that SETD2 drives lipotoxic damage in cardiometabolic HFpEF via transcriptional regulation of SREBP1 (**Fig. 8D**). Several lines of evidence support our conclusions: i) SETD2 is upregulated in cHFpEF and its chromatin mark H3k36me3 promotes the transcription of SREBP1, leading to lipid accumulation, lipotoxic damage and defective autophagy; ii) cHFpEF mice with CM-specific deletion of SETD2 do not develop LV hypertrophy, diastolic dysfunction and lung congestion as compared to SETD2^fl/fl^ cHFpEF littermates; iii) SETD2 deletion rescues detrimental changes of the myocardial lipidome such as TG and ceramide accumulation; iv) SETD2 depletion prevents lipid accumulation in cultured CMs exposed to metabolic stress while its overexpression mimics lipotoxic injury and defective autophagy; v) SREBP1 is required for the detrimental action of SETD2 in CMs; vi) SETD2 is upregulated in LV specimens from patients with cHFpEF vs control donors and positively correlates with cardiomyocyte stiffness, a main pathological feature of HFpEF; vii) pharmacological inhibition of SETD2 attenuates passive stiffness in CMs isolated from cHFpEF patients.

Cardiometabolic HFpEF is the most prevalent HFpEF phenotype and its incidence is expected to increase exponentially in decades to come due to the surge in obesity, metabolic syndrome, and hypertension in the general population.^2, 29–31^ Over 80% of patients with HFpEF are either overweight or obese, and in the recent STEP-HFpEF trial, the average BMI in HFpEF subjects exceeded 36 kg/m^2^.^32^ While the last decade brought significant advances in the treatment of several types of heart failure, only few drugs have shown an impact on HFpEF-related outcomes, with a consistent residual risk and a 5-year mortality rate ranging from 53% to 74%.^33^ Taken together, these data indicate that cHFpEF remains a poorly understood disease phenotype and new mechanism-based strategies are warranted.

Albeit a consistent body of evidence supports the notion that genes influence cardiometabolic features and outcomes, the "non-genetic regulation" of this process is gaining increasing attention. Plastic chemical changes of DNA/histone complexes - known as epigenetic changes - critically determine gene activity by rapidly modifying chromatin accessibility to transcription factors. Of interest, recent studies showed that changes in histone marks rather than CpG methylation underpin the transcriptional alterations observed in the failing heart.^11^ Indeed, epigenome-wide scale studies demonstrated that the combination of H3k36me3 and H3k27ac could explain more than 50% of disease-associated changes in gene expression in failing cardiomyocytes.^11^ SETD2 is the only histone methyltransferase responsible for H3k36me3 deposition in mammals.^34^ While the role of this methyltransferase has been thoroughly investigated in cancer and neurodevelopmental diseases, its role in the heart remains poorly understood. By performing bioinformatic analyses of available datasets, we show that H3k36me3 is a major orchestrator of the SREBP signaling pathway, a master regulator of lipid biosynthesis and uptake. Previous studies have confirmed that SREBP1 is upregulated in obese patients and mouse models of obesity and type 2 diabetes.^35, 36^ Consistently, SREBP1 has been shown to positively regulate the peroxisome proliferator-activated receptor-gamma (PPARγ), a key orchestrator of adipogenesis promoting lipid accumulation and insulin resistance.^38^ Indeed, upregulation of PPARγ has been shown to induce metabolic cardiomyopathy, a step preceding cardiometabolic HFpEF, by promoting transcription of several genes involved in lipid uptake, hydrolysis, and storage (i.e., *Cd36, Fabp3, Fabp4, Fas, Plin5, Lpl*).^27, 37, 38^ Previous ChIP-deep sequencing studies have already identified epigenetic modifications as central modulators of lipid metabolism, while pharmacological inhibition of the p300 histone acetyltransferase was shown to decrease the expression of key lipogenic genes.^39^

Accumulating evidence points to the fact that metabolic cardiomyopathy development is mainly the result of increased accumulation of triglycerides and ceramides in the heart.^39^ The mismatch between uptake and oxidation of fatty acids is gaining importance as a potent determinant of cardiac damage mainly via accumulation of free radicals and enhanced apoptosis.^40^ Interestingly, a large proportion of HFpEF patients displays increased TG accumulation and alterations in gene expression associated with contractile dysfunction.^41^ Using MS-based lipidomics, we demonstrated that SETD2 deletion in cardiomyocytes was able to reduce the levels of intracellular TGs as well as the synthesis of pro-apoptotic ceramides in the HFpEF myocardium, namely Cer[(d18:1/18:0)] and Cer[(d18:1/24:0)]. The importance of this result is supported by the notion that both these ceramides predict incident heart failure and mortality in patients with cardiovascular disease.^42^ We did not observe a significant effect of CM-specific SETD2 deletion on plasma lipids species. This result indicates that early alterations of the lipid landscape in cardiomyocytes represent an important event preceding lipotoxic injury and cardiomyocyte damage in the setting of HFpEF.

Autophagy plays a fundamental role in maintaining homeostasis and cellular function in the heart.^43^ Several studies demonstrated that autophagy protects against the development of pressure overload-induced or post-ischemic heart failure in animal models.^44, 45^ In our study, we show that a mouse model of cHFpEF displays increased lipid accumulation (both TGc and ceramides) and defective autophagy. Of interest, we found that SETD2 deletion restored the autophagic flux in HFpEF conditions leading to a decrease in lipotoxic stress and apoptosis. In our model, we show that SETD2-driven lipid accumulation suppresses cardiac autophagy while SETD2 deletion rewires the cardiac lipidome eventually leading to a restoration of autophagic flux and decreased cardiomyocyte apoptosis. In line with our data, previous work has shown a pivotal role of autophagy in cardiometabolic disease and obesity cardiomyopathy.^48–50^ Our results contribute to elucidate the link between metabolic stress, lipid overload and autophagy in the heart and demonstrate that specific chromatin marks, namely H3K36me3, have the ability to rescue myocardial lipotoxic injury thus leading to a restoration of autophagy and prevention of HFpEF-related cardiac dysfunction.

In recent years, a growing number of epi-drugs able to modulate chromatin remodeling has been developed. Of note, several of these compounds have already been approved by the FDA for the treatment of cancer, neurological and cardiovascular diseases.^46^ A recent study reported a highly specific, non-nucleoside SETD2 inhibitor for the treatment of acute myeloid leukemia. The identified inhibitor, known as Compound C13, effectively blocked the proliferation of two acute myeloid leukemia cell lines and led to decreased H3K36me3 levels.^47^ Furthermore, a screening campaign of the Epizyme proprietary histone methyltransferase-biased library identified EPZ-719 as an interesting compound for the investigation of SETD2 biology.^48^ Another study identified the small molecule EZM0414 as a first-in-class, selective, and orally bioavailable SETD2 inhibitor. Preliminary in vitro studies demonstrated that inhibition of SETD2 by EZM0414 suppresses proliferation in a panel of multiple myeloma (MM) and diffuse large B-cell lymphoma (DLBCL) cell lines.^49^ EZM0414 is currently being evaluated in a phase 1/1b open-label, multicenter, two-part study for the treatment of MM or DLBCL patients. In our study we tested the SETD2 inhibitor EZM0414 in cardiomyocytes obtained from LV biopsies of patients with cardiometabolic HFpEF. Cardiomyocytes from HFpEF patients displayed high levels of passive stiffness, a major feature of diastolic dysfunction and HFpEF. Of interest, pre-incubation of cardiomyocytes with EZM0414 was able to attenuate passive stiffness, suggesting that pharmacological editing of SETD2 results in functional improvement of human HFpEF cardiomyocytes. These findings could be explained by a SETD2-mediated reduction of lipotoxic injury eventually blunting cardiomyocyte oxidative stress and inflammation. The latter have been shown to play a prominent role in the regulation of cardiomyocyte stiffness by regulating titin functionality.^50^ Further work is needed to better elucidate the mechanisms underpinning the beneficial effects of SETD2 blockade on cardiomyocyte functionality in human HFpEF. In this perspective, our translational work set the stage for preclinical investigations addressing this important aspect.

Our study has some limitations. We cannot exclude that methylation of non-histone proteins by SETD2 plays a role in cHFpEF. However, we show that SETD2-driven H3k36me3 is highly enriched on the SREBP1 promoter and fosters lipid-related transcriptional programs eventually leading to cHFpEF features (myocardial lipid accumulation, LV hypertrophy, diastolic dysfunction and impaired exercise tolerance). Moreover, we also show that SREBP1 depletion in the presence of SETD2 overexpression prevents lipotoxic damage and defective autophagy. This set of experiments suggest that SETD2-mediated transcriptional regulation plays a major role in the setting of cHFpEF.

In conclusion, our translational study unveils a novel epigenetic mechanism and a potentially druggable target to prevent lipotoxic injury, defective autophagy and cardiomyocyte dysfunction in cHFpEF.

## Funding

This work was supported by the Swiss National Science Foundation (n. 310030_197557), the Swiss Heart Foundation (n. FF19045), the Olga Mayenfisch Foundation, the Swiss Life Foundation, the Kurt und Senta-Hermann Stiftung, the EMDO Stiftung, and the Schweizerische Diabetes-Stiftung (to F.P.); the Holcim Foundation and the Swiss Heart Foundation (to SC). S.A. and S.A.M. are the recipients of a Forschungskredit Candoc grant from the University of Zürich. This work was supported by EU’s Horizon 2020 research and innovation program under grant agreement No. 739593 to NH; DFG (Deutsche Forschungsgemeinschaft) HA 7512/2-4 and HA 7512/2-1 to NH.

## Author contributions

F.P. conceived the original hypothesis, planned the experiments, handled funding and supervision. F.P. and S.C. supervised the project. S.A., S.C, S.A.M., E.G., A.M. and M.H. carried out the core of the experiments. S.A and S.C. carried out all the in vitro experiments and molecular analyses in mice, performed statistical analyses, and worked on illustration of the data. S.A, S.C. and S.A.M performed HFpEF experiments and mouse harvesting; M.H. and N.H. measured passive stiffness in skinned CMs from cHFpEF patients and control donors. G.K. and T.H. performed lipidomic analysis in mouse hearts. L.V.H. provided myocardial samples from HFpEF patients and control donors. O.D., N.H. and F.R. provided critical intellectual feedback during manuscript preparation. S.A., S.C, and F.P. wrote the manuscript.

## Competing interests

The authors declare that the research was conducted in the absence of any commercial or financial relationships that could be construed as a potential conflict of interest.

## Notes

### Competing Interest Statement

The authors have declared no competing interest.

